# Nanobactericides Derived from Cinnamon Bark Extract: Phytochemical Profiling and Antibacterial Efficacy Against Bacterial Panicle Blight in Rice

**DOI:** 10.1101/2024.12.04.626863

**Authors:** Qamar Mohammed Naji, Dzarifah Mohamed Zulperi, Khairulmazmi Ahmad, Erneeza Mohd Hata

## Abstract

Bacterial panicle blight (BPB), caused by the Gram-negative aerobic bacterium *Burkholderia glumae (B.glumae)*, poses a significant threat to global rice production. Cinnamon bark extract (CBE) has demonstrated potent antioxidant and antimicrobial properties due to its high concentration of bioactive compounds, including eugenol and cinnamaldehyde. To enhance the efficacy and stability of these volatile compounds, this study employed nanotechnology and encapsulation techniques. The objective was to develop a CBE-based nanoformulation to inhibit *B. glumae* and control BPB in rice. CBE-chitosan (CBE-CS) nanoformulations were synthesized using ionic cross-linking between chitosan and trisodium phosphate (TPP) at various concentrations (0%, 0.5%, 1%, 2%, and 4% TPP). More than 15 active compounds were identified in CBE, including (Z)-3-Phenylacrylaldehyde, 2-Propenoic acid, 3-(2-hydroxyphenyl), cinnamaldehyde dimethyl acetal, and hexadecanoic acid. Bacterial membrane damage was significantly greater in treatments with CBE compared to untreated controls. The synthesized nanoparticles ranged in size from 43.66 nm to 106.1 nm, with encapsulation efficiencies between 48.65% and 48.78%, and loading capacities between 25.65% and 33.9%. Scanning electron microscopy (SEM) revealed spherical and homogeneous nanoparticles, while FTIR and XRD analysis confirmed the successful encapsulation of CBE in the chitosan nanoparticles. The antibacterial activity of the nanoformulations showed inhibition zones ranging from 7.5 to 11.8 mm, with the CBE-CS formulation containing 0.5% TPP demonstrating the highest efficacy (MIC = 15.6 μmol/ml; MBC = 31.25 μmol/ml).

## 1. Introduction

Since the dawn of civilization, archaeological evidence suggests that rice (*Oryza sativa*) has been a fundamental food source for humans, with its cultivation dating back to 1500-1000 BC (Verma & Srivastav, 2020). Today, rice remains a staple food for approximately 40% of the global population, particularly in the least developed countries (Zhang et al., 2022). The optimal climatic conditions for rice cultivation, including high humidity and abundant water, are prevalent in many Asian countries, especially in tropical regions (Rashid et al., 2022). In Malaysia, rice is the third most important crop after rubber and oil palm, with the states of Kedah and Perlis being key centers of rice production. However, rice fields are highly susceptible to a variety of diseases caused by microbes, fungi, and bacteria, which can severely affect productivity. Over 70 different diseases have been identified as impacting rice fields, with five to six major bacterial diseases attacking various parts of the rice plant, including the seedlings, leaves, leaf sheath, grains, stem, and roots (Faizal Azizi & Lau, 2022). One of the most severe bacterial diseases affecting rice worldwide, including in Peninsular Malaysia, is bacterial panicle blight (BPB), caused by *Burkholderia glumae* (*B. glumae*) (Ramachandran et al., 2021). This pathogen can cause up to a 75% yield loss and significantly reduce the quality of the affected plants, presenting a major challenge to rice cultivation. *B. glumae* thrives at optimal temperatures around 30°C, though it can survive in temperatures as high as 41°C. The bacteria infect seeds, enter plumules through stomata and wounds, and proliferate in the intercellular spaces of parenchyma during seed germination. This growth leads to the production of toxic substances like toxoflavin, which results in rice seedling rot (Rahman et al., 2024). Several strategies have been employed to manage BPB in rice cultivation (Chompa et al., 2022). Biological control methods, such as using harmless *Burkholderia* isolates, have been effective in reducing bacterial toxins and curbing rice grain rot.

Cultural practices, including the use of pathogen-free seeds, also help limit bacterial transmission (Kumar et al., 2023). While chemical treatments like oxolinic acid have been used to control seedling and inflorescence rot (Han et al., 2021), the rise of resistant *B. glumae* strains and the negative residual effects of chemicals on soil and plants have decreased their use. Recently, natural plant extracts such as cinnamon bark have emerged as promising antibacterial agents for BPB control (Sharifi-Rad et al., 2021). Cinnamon, a widely used spice, is derived from the inner bark of trees belonging to the genus *Cinnamomum* (Farooq et al., 2023). Both the bark and leaves are used in cooking and in various natural medicinal applications. Cinnamon bark extract (CBE) is rich in bioactive compounds such as eugenol, cinnamaldehyde, cinnamic acid, and coumarin (Mini Raj et al., 2023). These components exhibit diverse pharmacological properties, including antifungal, antibacterial, anti-inflammatory, antioxidant, antidiabetic, nematocidal, insecticidal, and anticancer effects (Anuranj et al., 2022).

With significant advancements in plant disease control and the advent of nanotechnology, organic nanoparticles (ONPs) and nanostructures have emerged as a promising field for managing plant diseases (Koul et al., 2021). While many methods exist for synthesizing inorganic nanoparticles from metals like gold and silver or semiconductors such as Si, ZnO, Ge, and GaAs, fewer techniques are available for producing ONPs. Organic nanomaterials have gained attention due to their unique structural and optical properties, which differ from their bulk forms. One of the most promising polymers for delivering agrochemicals and micronutrients in nanoparticles is chitosan (CS) (Mujtaba et al., 2020). Chitosan is a natural amino polysaccharide derived from fungal cell walls, the exoskeletons of insects and crustaceans, and other natural sources. Commercially, CS is produced by partially de-N-acetylating chitin. It is considered more effective than chitin due to its higher content of chelating amino groups and greater chemical modifiability.

As a cationic polyelectrolyte, chitosan exhibits unique properties, including its abundance, biocompatibility, biodegradability, and nontoxicity (Ul-Islam et al., 2024). Additionally, CS nanoparticles (NPs) offer great potential as nanocarriers, capable of encapsulating both hydrophilic and hydrophobic compounds (Pathak et al., 2023). In this study introduces CBE as a novel natural antibacterial ingredient against *B. glumae*. Compared to synthetic bactericides, the CBE nanoparticle formulation offers a superior strategy for combating plant pathogen infections in rice crops. The primary objective of this study is to develop a chitosan-loaded CBE nanoparticle formulation that effectively inhibits the growth of *B. glumae* in rice plants.

## 2. Materials and method

### 2.1. Raw materials

Chitosan (molecular weight 200 kDa, with a deacetylation degree of ≥75%) was procured for the study. Cinnamon bark (*Cinnamomum cassia*) was sourced from local markets in Malaysia. The raw materials were carefully transported to the laboratories of the Department of Plant Protection, Faculty of Agriculture, Universiti Putra Malaysia UPM, where they were utilized for experimental procedures and subsequent evaluation of results.

### 2.2 Extraction of Cinnamon Bark Extract

Cinnamon (Cinnamomum cassia) bark was obtained from market, wash, dry and ground into a fine powder using a blender. The powdered cinnamon (20 g) was then macerated in 200 mL of methanol (70% v/v) in a tightly sealed container for 48 hours at room temperature with intermittent shaking. After maceration, the cinnamon-methanol mixture was filtered through Whatman filter paper (Grade 1). The filtrate was concentrated under reduced pressure using a rotary evaporator at 40°C to remove the methanol and obtain the concentrated cinnamon extract.

### 2.3 Determination of the phytochemical compounds

A gas chromatography/mass spectrometry (GC-MS) analysis was performed to evaluate the volatile compounds and their abundances in CBE. This investigation using a Shimadzu QP-2010 Ultra GC-MS system as described by Wang et al. in (Wang et al., 2020). The system features a gas chromatograph connected to a mass spectrometer and utilizes an SLB-5ms capillary column (30 m length x 0.25 mm ID x 0.25 µm film thickness). The oven temperature program began at 50°C, then increased by 10°C per minute to 250°C, and finally increased to 300°C. The analysis was conducted at 70 eV, with a scan interval of 0.1 seconds, recording fragments ranging from 40 to 700 Da.

### 2.4. Preparation of cinnamon bark extract-loaded chitosan submicron emulsions

A nanobacterial bactericide targeting *B. glumae* was prepared using chitosan loaded with CBE, following the method described by (Maluin et al., 2019) with necessary modifications. The CBE was incorporated into chitosan nanoparticles using ionic gelation technique. Chitosan powder, sourced from Sigma Aldrich (low molecular weight, ≥ 75% deacetylation), was dispersed in a diluted acetic acid solution at a 1% (v/v) ratio and continuously stirred overnight at room temperature to form a 1% (w/v) CS solution. Subsequently, the CBE was mixed with Tween 80 (HLB 15.9) in a 1:1 (v/v) ratio, creating a CBE-Tween 80 mixture. This was stirred continuously for 1 hour at room temperature until a homogeneous solution was achieved. To initiate cross-linking, a modified ionic gelation method was employed. A 5 ml portion of the 1% CS solution was combined with the CBE-Tween 80 mixture under continuous stirring until the mixture became viscous. TPP solutions at concentrations of 0%, 0.5%, 1%, 2%, and 4% (w/v) were then added to 15 ml of the CS mixture, stirred at 1000 rpm for 15 minutes at room temperature. Finally, 5 ml of distilled water was added under magnetic stirring at room temperature and left for 24 hours to complete the formulation. The resulting nanobactericidal formulation was designated CBE-CS. Fig. 1 illustrates the preparation process of CBE-CS.

**Fig. 1:**
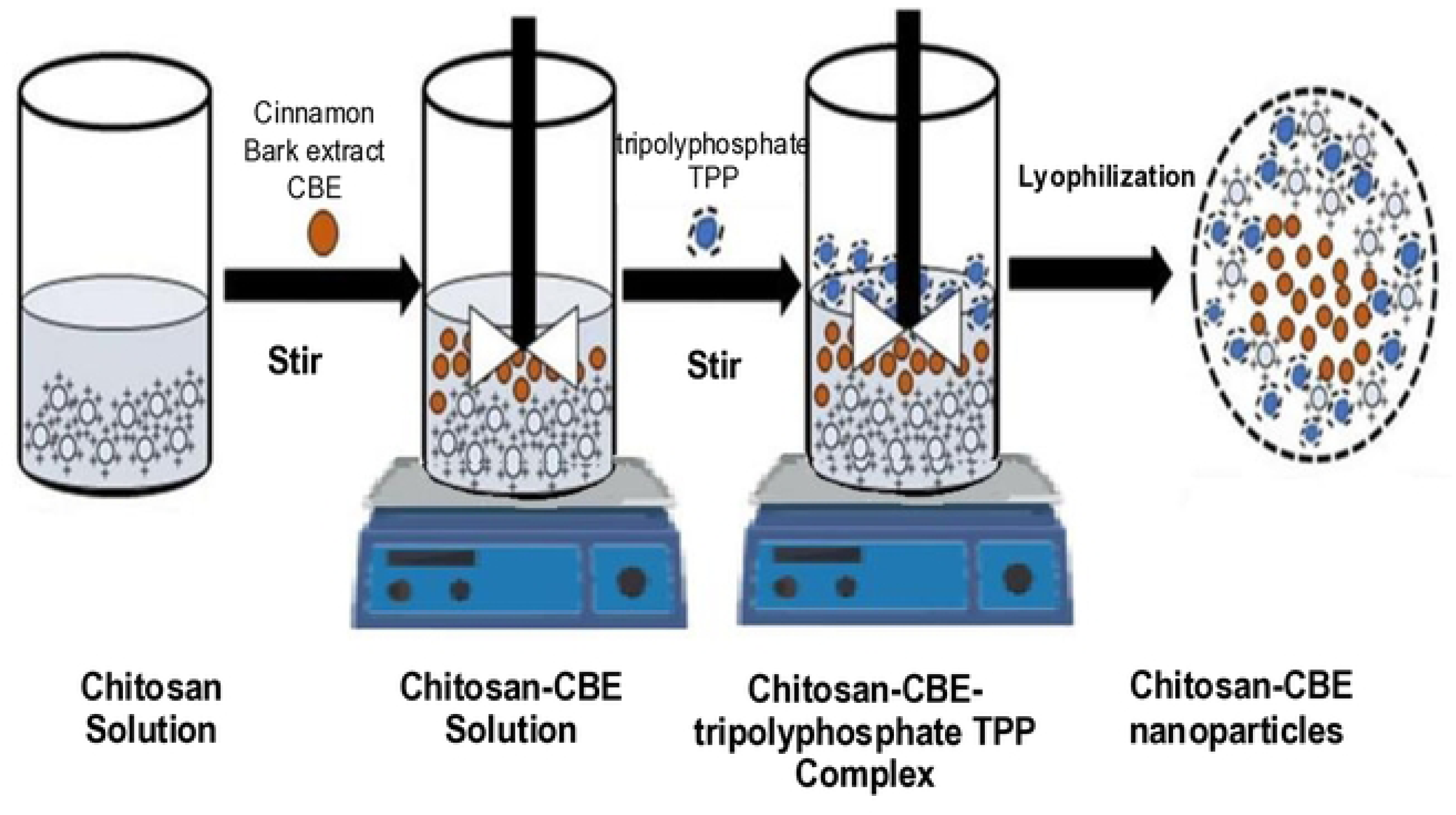
**Preparation of chitosan nanoparticles loaded with cinnamon bark extract using ionic gelation method**

### 2.5. Characterization of emulsions

#### 2.5.1. Particle size, polydispersity index and zeta potential

The mean particle size (PS), polydispersity index (PDI), and zeta potential (ZP) of CBE-CS nanoparticles were evaluated using a Zetasizer Nano ZS90 (Malvern, UK) through dynamic light scattering (DLS) at 25°C (Lunardi et al., 2021). Zeta potential, determined from electrophoretic mobility using the Helmholtz–Smoluchowski equation under a 40 V/cm electric field, assessed nanoparticle stability by indicating potential variation between the dispersion medium and the stationary fluid layer (Ağardan, 2020). Morphology was examined via SEM after coating the samples with a thin carbon layer. Stability was tested by storing various formulations at room temperature. All measurements, conducted in triplicate (n = 3), utilized water’s viscosity, refractive index, and absorption parameters provided by the Malvern software

#### 2.5.2. Transmission electron microscopy (TEM)

High-resolution TEM was employed to investigate the morphology of CBE-CS nanoparticles, with modifications to the method described by (Alghuthaymi et al., 2021). The samples were first diluted with distilled water and then deposited onto 200-mesh Formvar-coated copper grids for TEM imaging. The grids were subjected to low pressure and allowed to dry overnight at room temperature. The nanoparticles were observed under TEM without any staining.

#### 2.5.3. Fourier transform infrared spectroscopy (FTIR)

To analyze the active compounds in the CBE-CS nano extract, Fourier Transform Infrared (FTIR) spectroscopy was employed. This technique was utilized to identify the various chemical bonds within the molecules by generating an IR absorption spectrum (Veerasingam et al., 2021). The materials analyzed included powdered chitosan and TPP, along with four liquid samples of the CBE-CS nano extract and one sample of a simple CBE formulation for comparison.

#### 2.5.4. Encapsulation efficiency and loading capacity

Encapsulation efficiency (EE) and loading capacity (LC) of CB-CS nanobactericide formulations were evaluated using UV-Vis spectrophotometry with slight modifications to a standard protocol (Hoang et al., 2022). Freeze-dried nanobactericide (400 mg) was mixed with 5 mL of 1 M HCl, vortexed, and combined with 2 mL of ethanol, then incubated at 60°C for 12 hours. After centrifugation at 6000 rpm for 10 minutes at 25°C, the supernatant was analyzed with a UV-Vis spectrophotometer (200–400 nm). A calibration curve (R² = 0.9859) was used to quantify CBE. Triplicate measurements ensured accuracy, and EE and LC were calculated using standard formulas (Hoang et al., 2022).

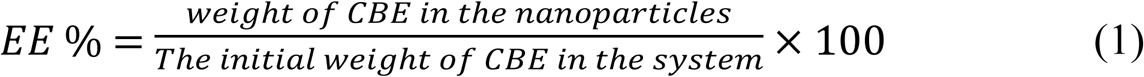

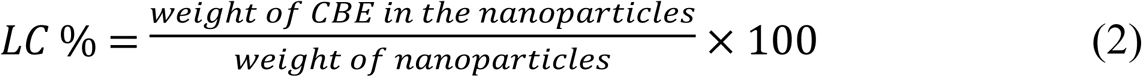

#### 2.5.5. Invitro release of nanobactericde formulation

The release analysis of the CBE-CS nanopesticide formulation was performed using a modified method from (Kaboudi et al., 2023). A 300 μL sample of the formulation was dissolved in 30 mL of phosphate-buffered saline (PBS, pH 7.4) at room temperature. At specific intervals, 3 mL samples were collected for analysis, replacing the same volume with fresh PBS to maintain consistency. The released CBE concentration was measured using UV-Vis spectroscopy, and the total release was calculated using formula (3).

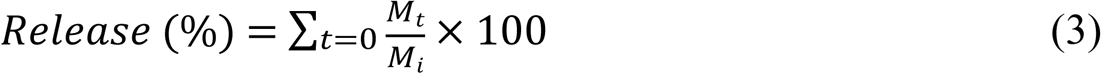

where *M_t_* represents the cumulative amount of CBE collected at time t, and *M_i_* is the initial weight of CBE incorporated into the formulation. This formula provides the percentage of CBE released over time, reflecting the release kinetics and efficiency of the nanopesticide formulation.

#### 2.5.6. Evaluation of the antibacterial activity of CBE-CS

The antibacterial efficacy of the nano-CBE-CS formulation against *B. glumae* was evaluated using disk diffusion assays. A *B. glumae* suspension (10⁸ CFU/mL) was prepared in sterile distilled water and spread evenly on Mueller-Hinton Agar (MHA) plates. Sterile Whatman filter paper discs (6 mm diameter) were impregnated with 20 μL of formulations (CBE-CS 0, 0.5, 1, 2, and 4) in a laminar flow hood, with streptomycin (15 μg/mL) and distilled water serving as positive and negative controls, respectively. The discs were placed on the inoculated plates, left at room temperature for 30 minutes for diffusion, and incubated at 37°C for 24 hours. Inhibition zones were measured in millimeters, and bacterial inhibition percentages were calculated using formula (4). Each experiment was performed in triplicate and repeated twice (Chavez-Esquivel et al., 2021).

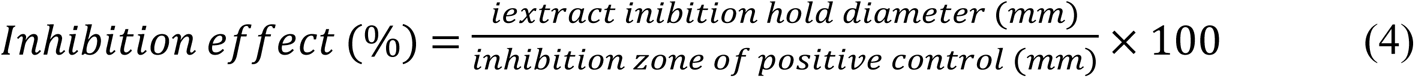

MEC and MBC tests were performed by double dilution method on the CBE-CS nanobactericidal formulation at a concentration of 0.5% TPP due to the small diameter of the nanoparticles and the high zeta potential value of this concentration.

#### 2.5.7 Time killing analysis

To evaluate the bactericidal effects of the CBE-CS nanocide formulation, a growth curve analysis was conducted. In this study, a 96-well microplate was used to assess the bactericidal activity of the CBE-CS nanocide formulation. Each well was first filled with 150 μL of Mueller-Hinton Broth (MHB), followed by the addition of 150 μL of the CBE-CS nanocide formulation at concentrations corresponding to 0.5 MIC, 1 MIC, and 2 MIC. A control well received only 1% Tween 80. Subsequently, 50 μL of *B. glumae* suspension (10˄7 CFU/mL) was added to each well. The microplate was incubated at 37°C, and the optical density (OD) at 600 nm of the liquid culture was measured at regular intervals using a spectrophotometer to monitor bacterial growth (Cava-Roda et al., 2021)

## 3. Results and discussion

### 3.1. Determination of phytochemical constituents

The qualitative and quantitative analysis of CBE was conducted using GC-MS. Detailed on the various bioactive compounds found in CBE is presented in Table 1.

**Table 1:**
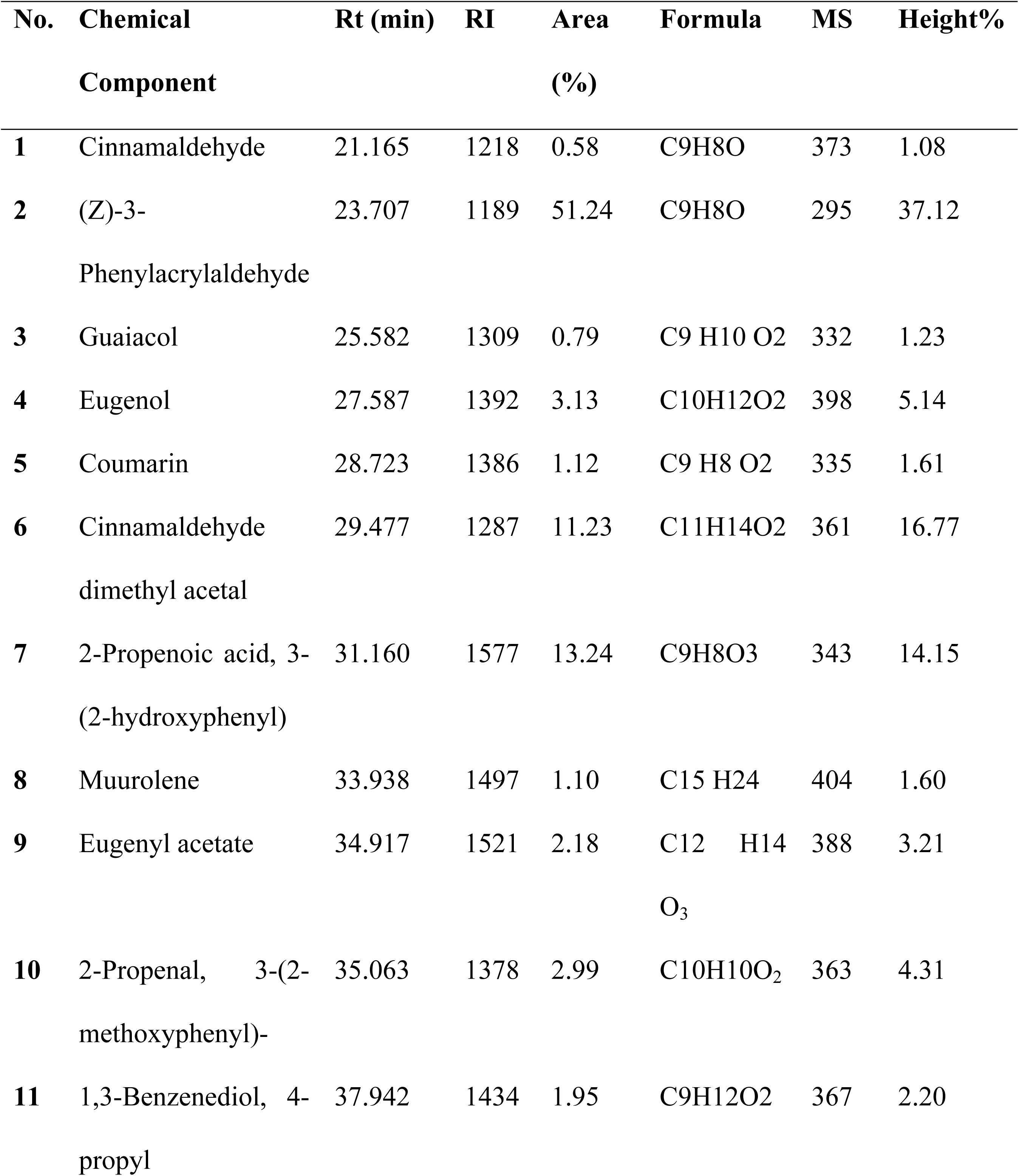
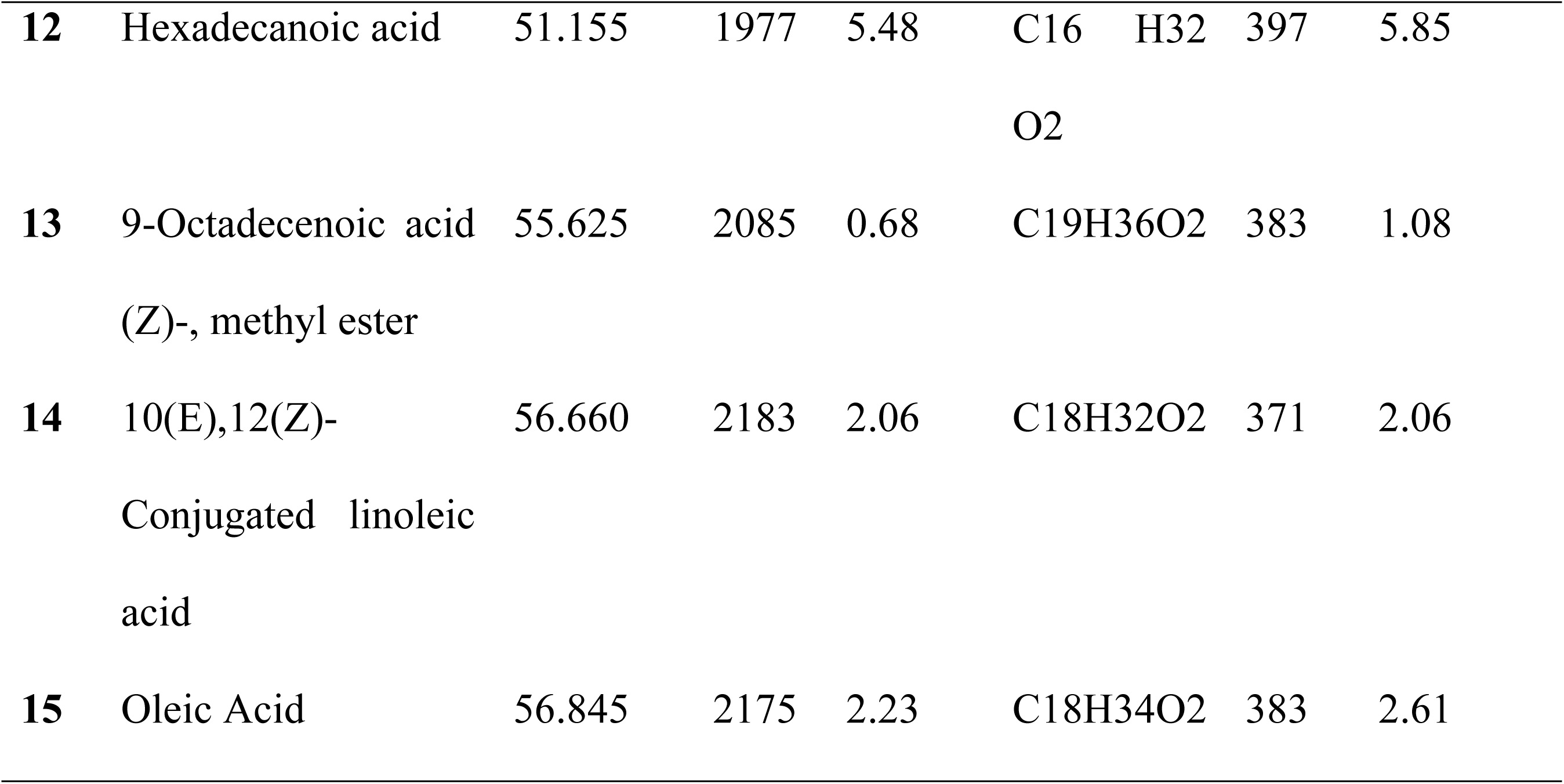
Phytochemical constituents in CBE identified by GC-MS, comparing Retention Times (Rt), Retention Index (RI), Area %, molecular formula, mass spectral (MS), and Height Data with databases FFNSC1.3.lib, NIST11.lib, and WILEY229.lib.

The identification of constituents in CBE was performed by comparing their retention times and retention indexes with data from libraries such as FFNSC1.3.lib, NIST11.lib, and WILEY229. The gas chromatography analysis of CBE revealed the presence of 15 active substances. Among these compounds, (Z)-3-Phenylacrylaldehyde was found to be the most predominant, constituting 51.24% of the total composition. Following this, 2-Propenoic acid, 3-(2-hydroxyphenyl) accounted for 13.24%, while Cinnamaldehyde dimethyl acetal and Hexadecanoic acid were present at concentrations of 11.23% and 5.48% respectively. Additionally, 2-Propenal, 3-(2-methoxyphenyl) was identified at 3.99%, followed by Eugenol (3.13%), Oleic Acid (2.23%), and Eugenyl acetate (2.18%). Furthermore, 10(E),12(Z)-Conjugated linoleic acid was detected at 2.06%. The analysis also revealed the presence of 4 aldehydes within the CBE. Apart from the compounds, the remaining constituents were found in smaller quantities, each comprising less than 2% of the specimen. Specifically, 1,3-Benzenediol, 4-propyl was identified at 1.95%, followed by Coumarin (1.12%), Muurolene (1.10%), and Guaiacol (0.79%). Additionally, 9-Octadecenoic acid (Z)-, methyl ester and Cinnamaldehyde were detected at concentrations of 0.68% and 0.58% respectively. Fig. 2 illustrates the chemical analysis and characterization of CBE conducted using GC-MS, highlighting the identification of its key compounds.

**Fig. 2:**
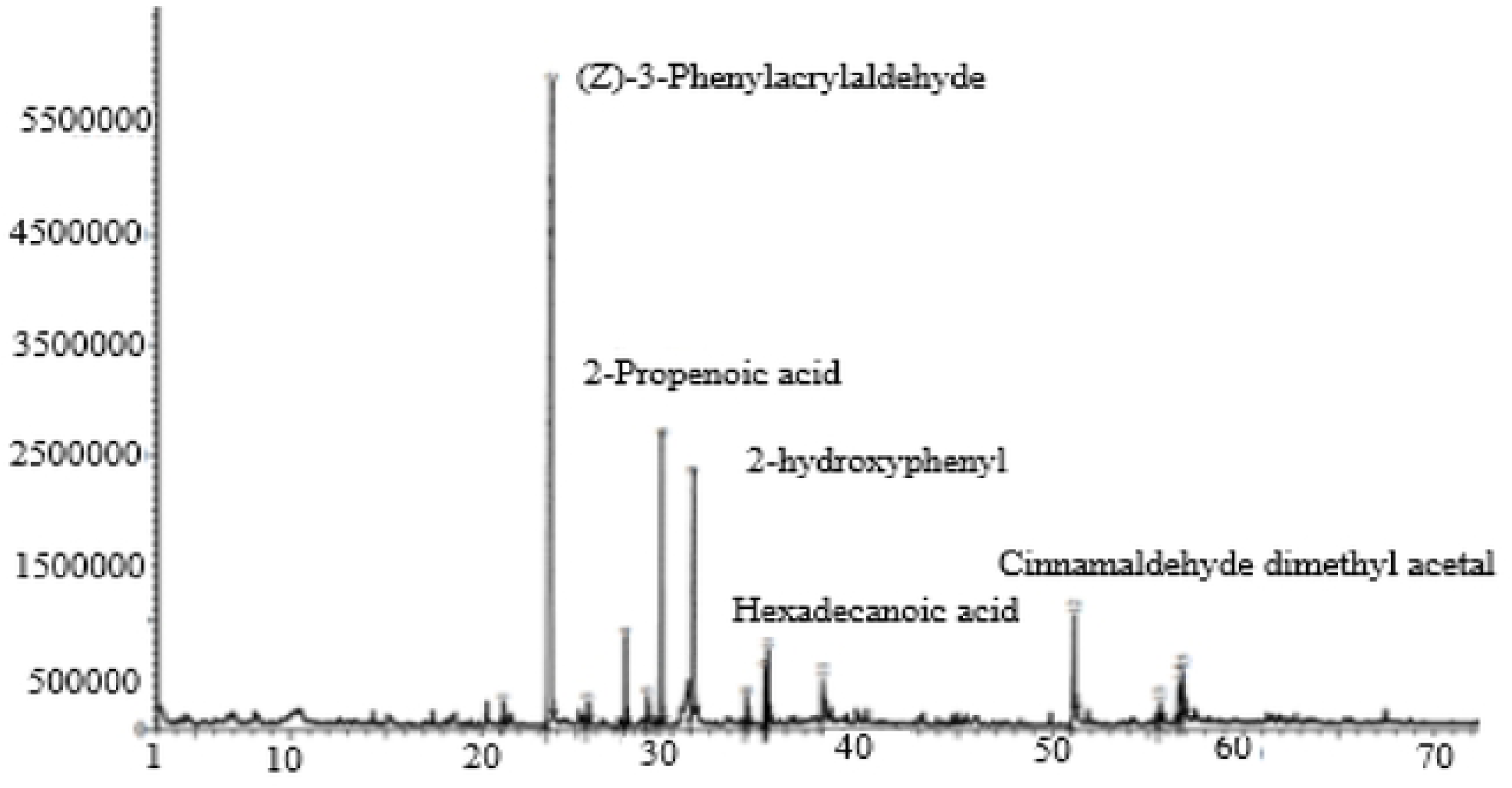
**GC-MS chromatogram of CBE, depicting phytochemical analysis characterization of CBE**

### 3.2. Size, polydispersity index and zeta potential

The physical properties of nanomaterials such as PZ, PDI, and ZP contribute to determining their behavior pattern within different biological environments. The size distribution was measured in a Zeta sizer Pro (Malvern Instruments Ltd, UK) and the results obtained represent the average value of three samples. While the zeta potential was measured in triplicate for the dispersed nanoparticles of each nanoparticle suspension using a Zeta sizer Pro with a DTS1070 zeta cell (Malvern Instruments Ltd, UK). From the results presented in Table 2, it is evident that the concentration of TPP plays a crucial role in determining the average hydrodynamic diameter of the particles in the CBE-CS.

**Table 2:** The impact of TPP concentration on the CBE-CS nanobactericide formulation’s average PS, PDI, and ZP.

| Nano bactericide | Average size (d. nm) | PDI | Zeta Potential (mV) | Mob $\mu\text{mcm/Vs}$ | Cond mS/cm |
| --- | --- | --- | --- | --- | --- |
| 0% TPP | 435.68±22.1 | 1.00±0.00 | 7.79±0.04a | -0.47±0.04d | 0.97±0.02a |
|  | 4a | a |  |  |  |
| 0.5% | 43.66±0.23a | 0.28±0.03 | 1.78±0.04b | 0.61±0.03a | 0.34±0.02c |
| TPP |  | c |  |  |  |
| 1% TPP | 51.72±0.04a | 0.57±0.02 | 1.01±0.01b | -0.20±0.06c | 0.90±0.03a |
|  |  | b |  |  |  |
| 2% TPP | 79.67±0.17a | 0.68±0.01 | -2.58±0.08c | 0.13±0.03b | 0.53±0.07b |
|  |  | b |  |  |  |
| 4% TPP | 106.06±0.42 | 0.96±0.00 | -6.11±0.61d | 0.07±0.01b | 0.90±0.02a |
|  | a | a |  |  |  |
| **Mean 1-** | 1.08E+04 | 0.661 | 0.396 | 0.03099 | 0.732 |
| **30** |  |  |  |  |  |
| **Std Dev** | 1.96E+04 | 0.325 | 4.81 | 0.3767 | 0.258 |

At 0% TPP, the particles exhibited a relatively large average size of 790.5 d.nm with a PDI index of 1, indicating significant polydispersity and potential instability in the formulation. As the TPP concentration was increased to 0.5%, there was a dramatic reduction in particle size to 43.66 d.nm, accompanied by a more uniform distribution, as reflected by a lower PDI of 0.288. This initial decrease in size suggests that TPP may facilitate the formation of more stable, smaller nanoparticles by promoting cross-linking within the extract. However, as the TPP concentration increased further, the average particle size began to rise, with measurements of 51.72 d.nm (PDI 0.481) at 1% TPP, 79.82 d.nm (PDI 0.576) at 2% TPP, and 106.1 d.nm (PDI 0.96) at 4% TPP.

This trend of increasing particle size with higher TPP concentrations indicates a shift towards agglomeration, likely due to excessive cross-linking or the potential leakage of TPP onto the particle surfaces, which promotes sedimentation and aggregation. The increasing PDI values with higher TPP concentrations further support the notion of growing heterogeneity within the particle population (Somenath Das et al., 2020) and (Hasheminejad et al., 2019).

Consistent with earlier research, particularly studies by Huang et al. in (Huang et al., 2009) and Calvo et al. in (Calvo et al., 1997). it was found that PDI values below 0.45 correspond to a narrow size distribution, like what was observed at 0.5% TPP in the current study. These previous studies demonstrated that the cross-linking of chitosan with TPP resulted in the formation of submicron particles with PDI values less than 1, which aligns with the findings of this investigation. In this study, the positive ZP values observed indicate a higher concentration of hydrogen ions (H+) relative to hydroxyl ions (OH−) around the particle surfaces. This phenomenon is typically attributed to the protonation of amino groups (NH3+) present in chitosan (Prasad et al., 2022). The observed decrease in zeta potential values with increasing TPP concentrations suggests a reduction in positive peripheral charges. This is likely due to the interaction of TPP with the particle surface, which diminishes the number of protonated amino groups (NH3+) on the chitosan particles (Fernández-Díaz et al., 2017). The study findings align with previous reports indicating that for an emulsion to remain physically stable, an electrical potential close to ±30 mV is required. Notably, all concentrations of the nano extract tested in the current experiment achieved this stability threshold. These results corroborate the findings of earlier studies, including those by (Deshi et al., 2024), which also reported similar behaviors in systems involving chitosan and zeta potential measurements.

### 3.3. Transmission electron microscopy (TEM)

The TEM images in Fig. 3 (A) revealed that the nanocomposite was nearly circular when TPP was absent. In contrast, Fig.3 (B to D) demonstrated that the average size of the nanocomposite increased to 31 nm, 60 nm, and 73 nm for CBE-CS 0.5, CBE-CS 1, and CBE-CS 2, respectively, with a sharp and distinctly crystalline surface. The results also indicated that particle size increased with rising TPP concentration, as observed in the TEM images. The spherical nanoparticles exhibited lighter edges than their centers, suggesting TPP coating. Despite some particle clustering, the overall composition displayed a homogeneous distribution of the extract, with a distinctly crystalline nanoparticle surface.

**Fig. 3:**
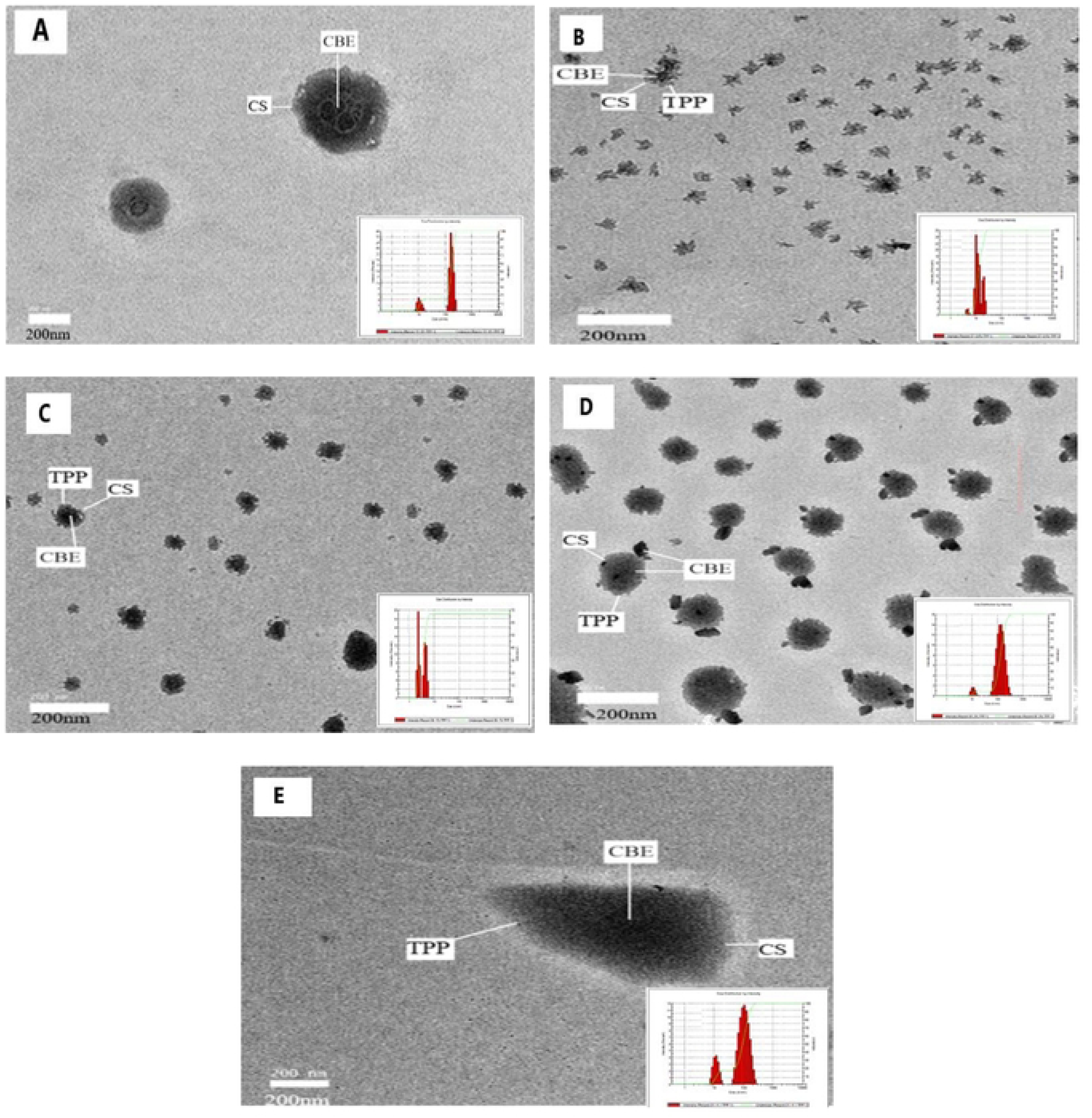
**TEM images of CBE-CS nanobactericide formulation with different concentrations of TPP: (A) 0%, (B) 0.5 % (C) 1%, (D) 2% and (E) 4% (scale bar is 200 nm)**

### 3.4 Morphology and FTIR spectroscopy of emulsions

The results of FTIR spectroscopy analysis used to determine the chemical composition of the samples are shown in Fig.4. The characteristic values of pure chitosan indicate the presence of many active chemical compounds, as the peak at 3556 cm-1 indicates the presence of O-H alcohol stretching, the peak at 3329 cm-1 indicates N-H stretching of the secondary amine, the peak at 2241 cm-1 indicates C≡N stretching, the peak at 1519 cm-1 indicates N-H bending, the peak at 1068 cm-1 indicates C-O stretching, while the peak at 732 cm-1 indicates C=C bending. In the FTIR spectra of TPP, the peak at 1127 cm⁻¹ corresponds to the O–P=O stretching vibration, while the peaks at 1211 cm⁻¹ and 1097 cm⁻¹ are associated with P=O and PO₃ stretching vibrations, respectively, indicative of phosphorylated structures. Other significant peaks include 1116, 1087, and 1028 cm⁻¹ (C–O, C–C, and C–H vibrations), all of which are crucial in characterizing the carbohydrate backbone. The infrared spectrum of cinnamon bark showed a sharp peak with high intensity at 1662 cm-1, which indicates the stretching of the aldehyde carbonyl group C=O, which represents a high concentration of aldehyde in CB. Also, the presence of significant peaks at the range of 3630 to 2735 cm-1 indicated the stretching of alcohol O–H, the peak at 1134 cm-1 indicated the stretching of C–O, and finally the peak at 759 cm-1 indicated the bending of C=C. The cross-linking of the chitosan polymer with TPP and CBE in the different concentrations of nano extract CBE-CS resulted in a significant change in the peak groups at the peak 3329 cm-1 related to amide (N-H stretching of secondary amine) and the peak at 3379 cm-1 related to N-H stretching of primary amine. It is also evident that the peak shape became distinctly sharp due to the enhancement of hydrogen bonding, indicating the participation of TPP and amine groups in chitosan in the chemical reaction. While the presence of new absorption peaks at 2870 cm-1 indicates O-H stretching and at 1624 cm-1 indicates C=C stretching, while the high intensity stretching band at 1053 cm-1 indicates C-O stretching. In the nano-extract CBE-CS at 0% of TPP, the distinct peaks at 1662 cm-1 indicated the extension of aldehyde carbonyl group by carbon = oxygen) and the peak at 1134 cm-1 indicated the extension of aldehyde carbonyl group. The results obtained in this study agree with the previous studies in references (Somenath Das et al., 2020), (Prasad et al., 2022), (Deepika et al., 2021), (Deepika et al., 2021), and (Chaudhari et al., 2022).

**Fig 4:**
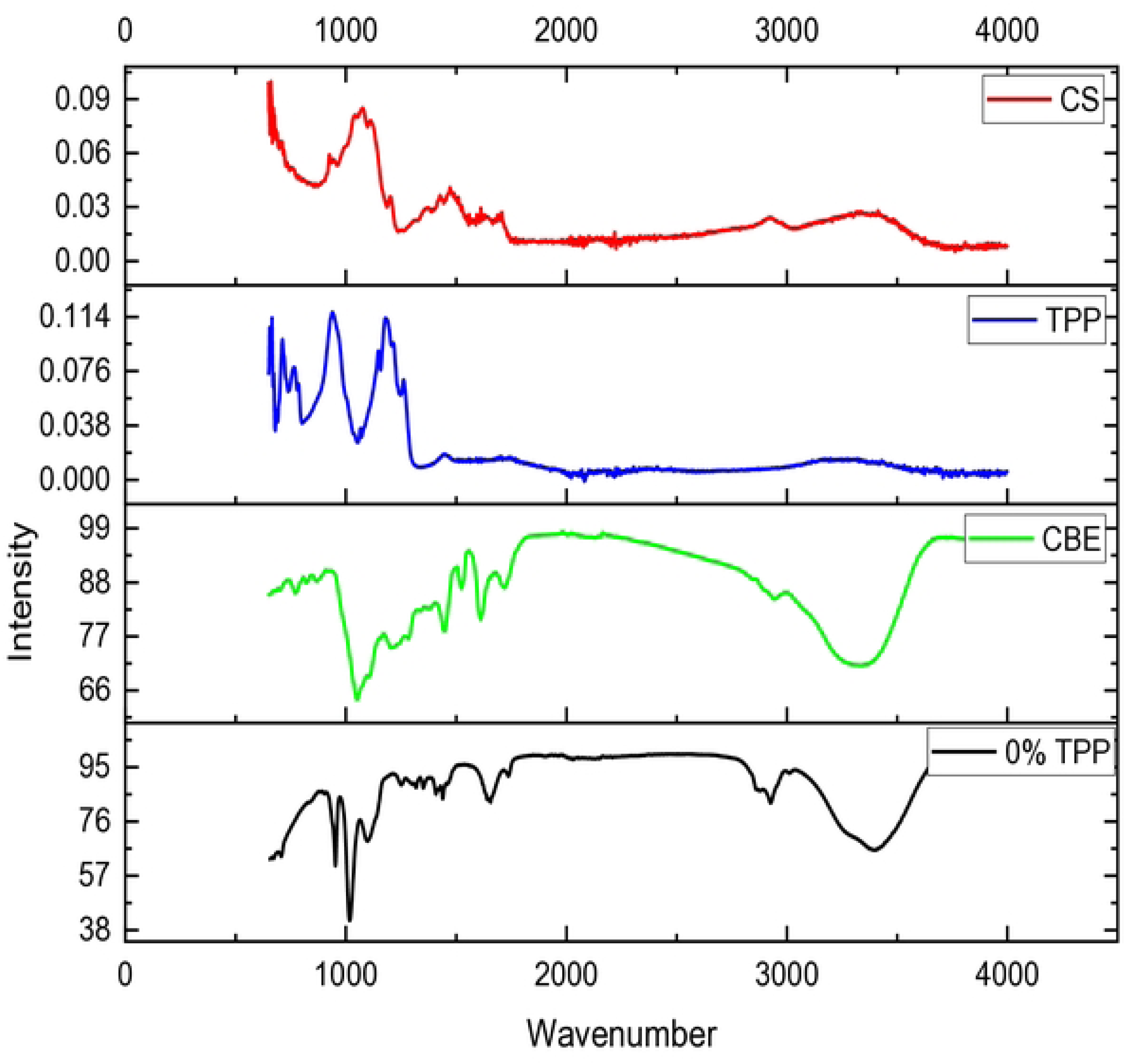
**FTIR spectra of chitosan, TPP, CBE, CBE-CS at 0% TPP**

### 3.5 Encapsulation efficiency (EE) and loading capacity (LC)

The EE and LC of CBE-CS nanocapsules were determined using UV-visible spectrophotometry at the maximum absorption wavelength of CBE-CS. The results, illustrated in Fig.5, demonstrate the impact of varying TPP concentrations on the LC and EE of the nanocapsules. As the TPP concentration in the nanoemulsion increased, there was a notable positive effect on the loading capacity of the nanocapsules. Specifically, the LC values rose from 25.65% to 33.9% as the TPP concentration increased from 0% to 4%. This trend indicates that higher TPP concentrations enhance the ability of the nanocapsules to incorporate CBE. However, if TPP is excessively increased, the nanoparticle structure might become too rigid and dense, which could reduce the available space for drug loading, thereby decreasing the loading capacity. On the other hand, the encapsulation efficiency remained relatively stable across all TPP concentrations, with values ranging between 48.65% and 48.78%. The consistent EE suggests that a saturation point may have been reached, beyond which further increases in TPP concentration do not significantly impact encapsulation. These findings are consistent with previous studies. For instance, (Culas et al., 2023), (Deepika et al., 2021), and (Amiri et al., 2021) also observed stable encapsulation efficiencies in similar formulations. Conversely, studies by (Prasad et al., 2022), and (Deepika et al., 2021) reported increases in both EE and LC with higher TPP concentrations.

**Fig. 5:**
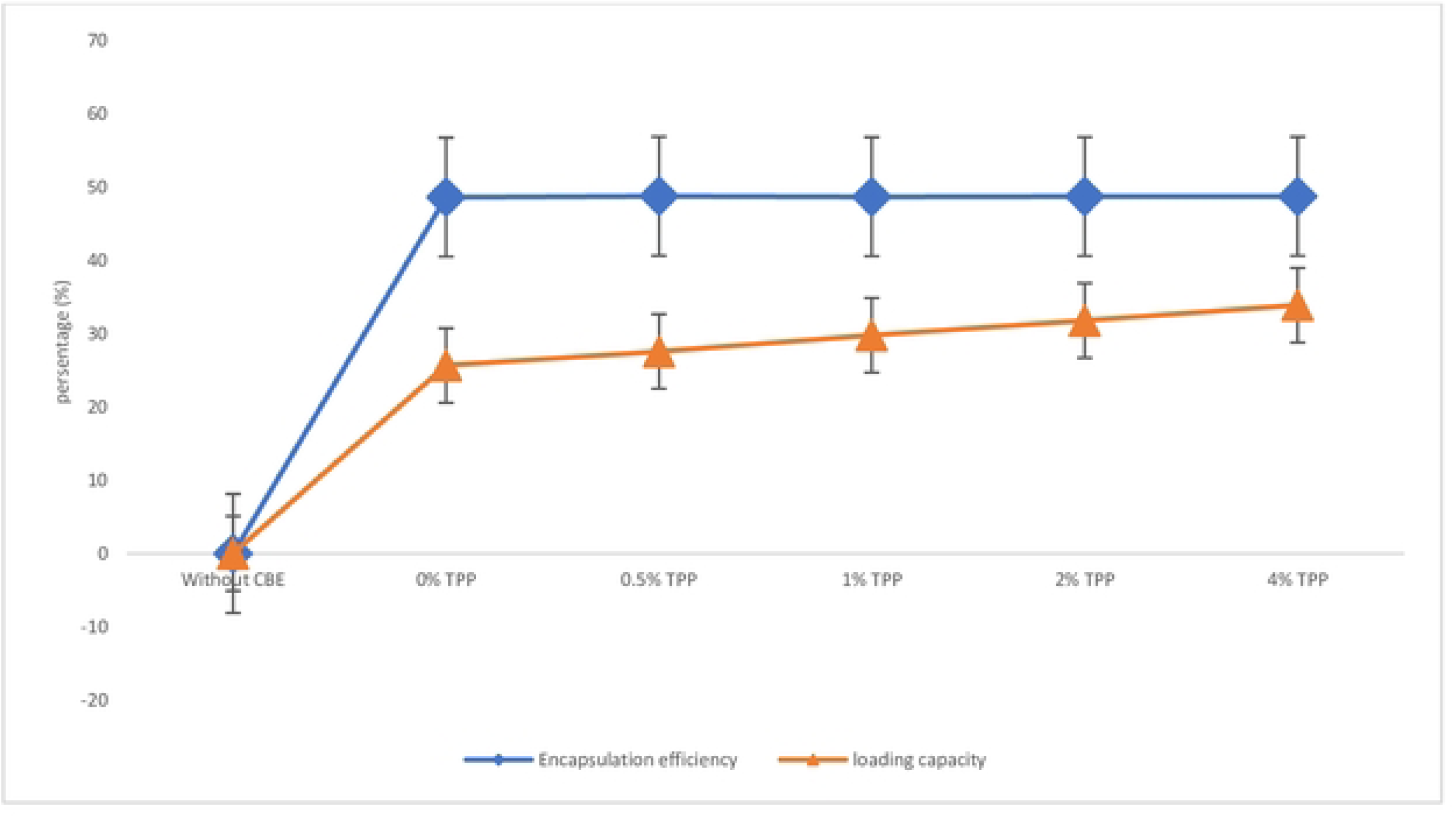
**Effect of CBE loading at different TPP concentrations on encapsulation efficiency and loading capacity (Value indicates mean of four replicates).**

### 3.6 In vitro release test

In vitro release kinetics test was performed where dried capsules were added in aqueous-alcoholic medium at a volume ratio of 1:1. UV-visible spectroscopy at 286 nm wavelength was used to evaluate the concentration of CBE (Ramasamy et al., 2017). Readings were taken at times t = 0 (starting time), (t) 10, 20, 30, 60, 240, 1440, and 2880 min for triplicates. Fig.6 shows the release of CB extract over time. As shown, after 1440 min, the release rate of CB extract was 0.44 μmol/mL. The curve is divided into two separate parts. The CBE concentration increases quickly in the first section, which corresponds to the test’s initial hours. In enclosed systems, this behavior is frequently seen (Amiri et al., 2021). As the CB contained in the capsule core slowly reaches the capsule surface, the second area of the curve shows a nearly constant CBE concentration, (Shetta et al., 2019) and (Sibaja et al., 2015). Based on the concentration difference, these data show that the CB release is connected to the concentration on the inside and external region of the capsule.

**Fig. 6:**
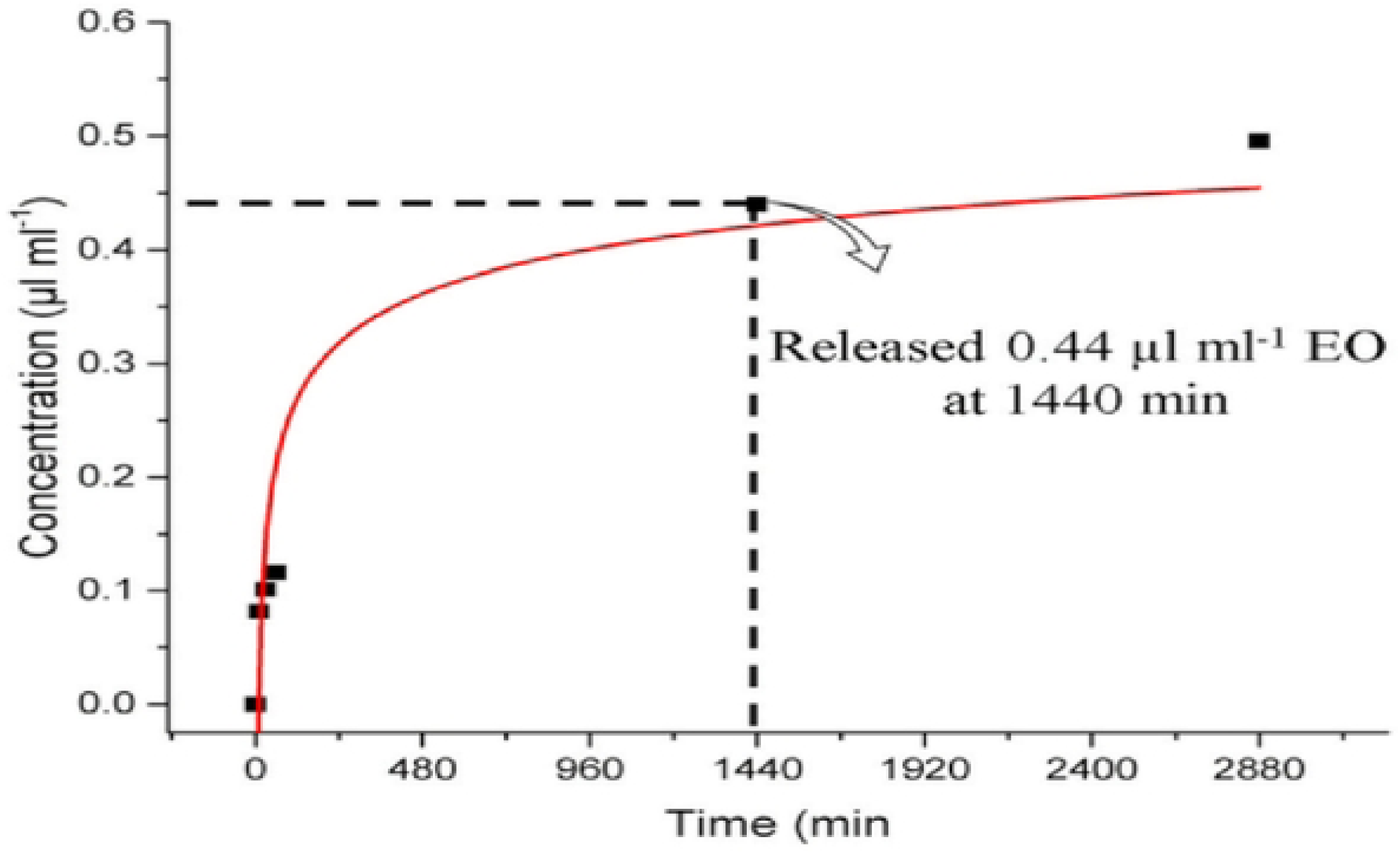
**The release kinetics of CB extract over time**

### 3.7. Assessment of antibacterial activity, MIC, and MBC

Inhibition zone formation, MIC, and MBC are critical measures of antimicrobial effectiveness. The in vitro evaluation of different CBE-CS nano-extract formulations against *B. glumae* demonstrated varying degrees of antibacterial efficacy, as measured by the DIZ, the percentage of inhibited radial growth (PIRG), and the increment of activity of the test discs. The results, summarized in Table 3, reveal that all nano-formulations containing different concentrations of TPP were effective, albeit to varying extents.

**Table 3:** Disc diffusion method-based antimicrobial activities of CBE-CS nanobactericide formulation and the antibiotic streptomycin against *B. glumae*.

|  | CBE-CS formulation |  |  |  |  | DH2O | Pos.<br>control |
| --- | --- | --- | --- | --- | --- | --- | --- |
|  | 0 % | 0.5 % | 1 % | 2% | 4% |  |  |
|  | TPP | TPP | TPP | TPP | TPP |  |  |
| **Inhibition zone** | 8.6 ± | 11.3 ± | 11.5 ± | 11.6 ± | 11.3 ± | 0.0 ± | 20.0 ± |
| **(mm)** | 0.33<sup>c</sup> | 0.33<sup>b</sup> | 0.28<sup>b</sup> | 0.76<sup>b</sup> | 0.33<sup>b</sup> | 0.0<sup>d</sup> | 0.00<sup>a</sup> |
| **PIRG** | 31.7 | 46.7 | 47.1 | 47.9 | 46. | - | - |
| **(%)** |  |  |  |  |  |  |  |
| **Effectiveness** | - | 47.4 | 0.8 | 1.72 | -2.5 | - | - |
| **Increase (%)** |  |  |  |  |  |  |  |

The data indicates that the nano-extract with 0% TPP concentration exhibited the lowest inhibition zone, measuring 7.8 mm. As TPP concentrations increased, a corresponding significant increase in DIZ was observed, with values ranging from 11.5 to 11.8 mm for concentrations between 0.5% and 2% (P>0.05). Notably, the nano-extract with 4% TPP concentration achieved a DIZ of 15.5 mm, while the positive control treated with streptomycin resulted in the highest DIZ at 24.6 mm, outperforming all other nano-pesticide formulations. Furthermore, the PIRG index showed a positive correlation with increasing TPP concentrations, indicating enhanced antibacterial activity. The most significant improvement in activity was observed at a TPP concentration of 0.5%, with a remarkable increment of 47.4% in effectiveness. When determining the MIC, the color change in the microtiter plate wells, which transitioned from colorless to bright red, indicated the presence of live bacteria and the varying impact of TPP concentrations on bacterial inhibition. The MIC value was determined to be 31.25 μmol/mL, while the MBC value was 15.6 μmol/mL. These findings underscore the potent antibacterial activity of the CBE-CS nano-extract against *B. glumae*, highlighting its potential as an effective antimicrobial agent.

### 3.8. Time-killing analysis

The time-killing test was conducted to assess the bactericidal efficacy of CBE-CS 0.5% TPP nano-extract at varying concentrations of MIC (0.5, 1, and 2) against *B. glumae*. A bacteria-free control was used to establish a baseline, with its optical density (OD) remaining steady after an initial increase during the first 18 hours of incubation. The results of the growth curve are illustrated in Fig.7. The nano-extract demonstrated a concentration-dependent effect on the growth rates of *B. glumae* throughout the 24-hour period. All tested concentrations of MIC showed a reduction in bacterial growth rates within the first 4 hours. For concentrations of 0.5 MIC and 1 MIC, a slight decrease in the growth rate was observed after the initial 4-hour. In contrast, the nano-extract at 2 MIC exhibited a more pronounced impact, with a significant decrease in the growth curve observed after the first 4 hours until 8 hours. This result indicates that all MIC concentration was highly effective, rapidly reducing the bacterial population and preventing further growth.

**Fig. 7:**
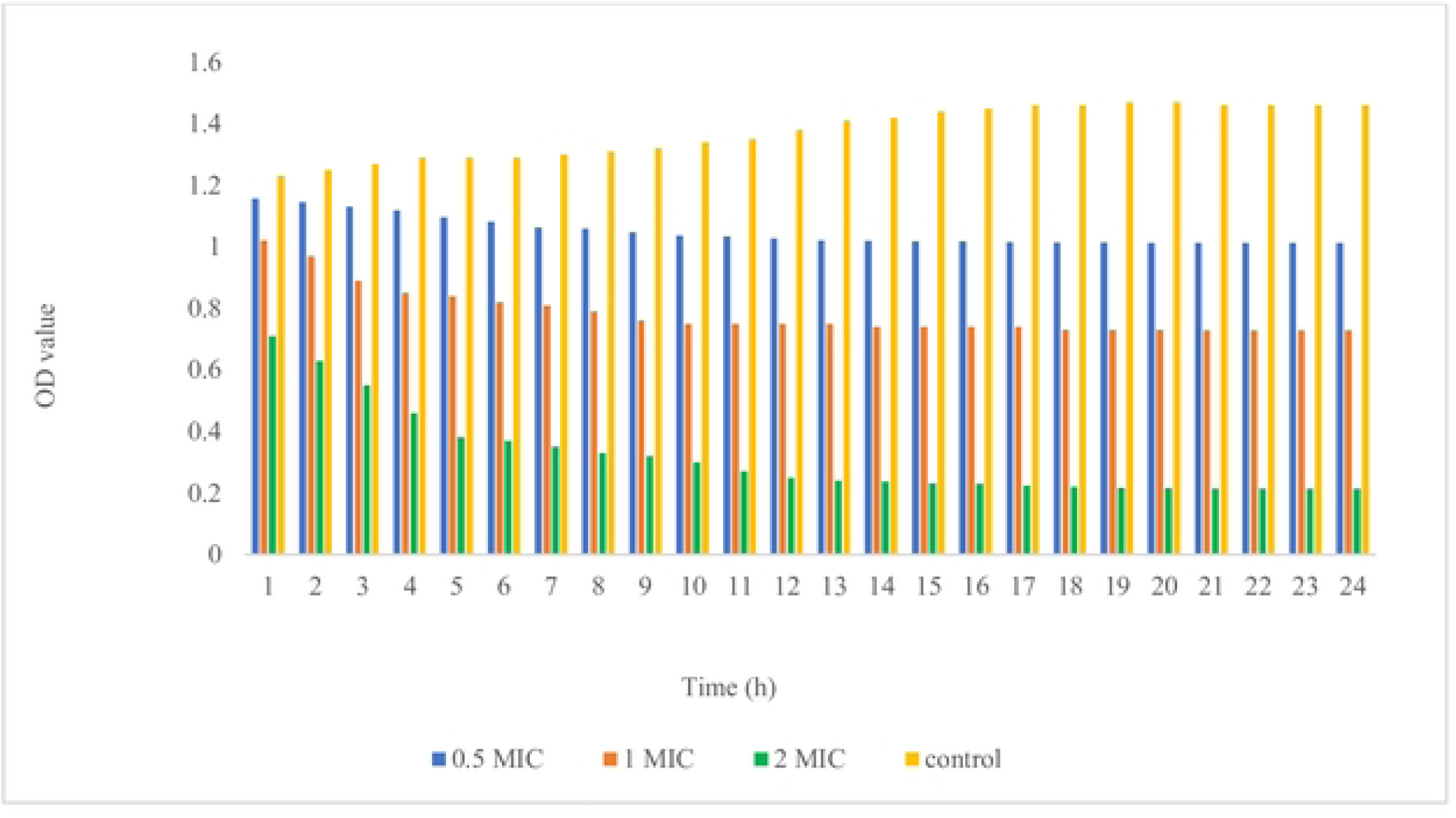
Time Killing analysis of the CBE-CS formulation against *B. glumae* at (0.5, 1, 2) of MIC. The absorbance OD at 600 nm was measured until the 24. **h.**

## 4. Conclusion

In this study, we developed a CBE-loaded nanoencapsulation aimed at controlling bacterial panicle blight caused by *Burkholderia glumae* in rice plants. The nanoencapsulation was formulated with varying concentrations of TPP (0%, 0.5%, 1%, 2%, and 4%) and characterized for its internal structure, morphology, and size using TEM. The results revealed that all nanoparticles were within the nanometer scale, with the introduction of TPP significantly influencing their morphology. Specifically, TPP contributed to a more spherical shape, likely due to the increased rigidity from cross-linking between TPP and chitosan (CS). Particle size analysis during dissolution in deionized water showed an inverse relationship between TPP concentration and nanoparticle size, with the smallest particles observed at 0.5% TPP. PDI analysis indicated a shift from monodisperse to polydisperse systems with higher TPP concentrations, suggesting decreased stability due to increased agglomeration. This instability may result from gel formation through inter-and intra-particle cross-linking. Zeta potential measurements revealed that increasing TPP concentrations caused a shift from positive to negative surface charges, attributed to the negatively charged functional groups of cinnamon molecules coating the nanoparticles. Higher zeta potential values indicate superior stability and functionality, while the observed negative zeta potentials may influence nanoparticle aggregation. FTIR analysis confirmed the successful encapsulation of CBE within the CS nanoparticles, as evidenced by characteristic absorption bands and peaks associated with both CBE and CS functional groups. The release of bioactive substances from the nanoparticles likely followed mechanisms such as disintegration, surface erosion, desorption, and diffusion. Initial rapid release within the first two hours, followed by a slower release, is crucial for effective bacterial control. The zone of inhibition tests confirmed the antibacterial efficacy of the CBE-CS nanobactericide against *B. glumae*, with a significant reduction in microbial count compared to the distilled water-negative control, where no inhibition was observed. This suggests that the CBE-CS formulation offers promising potential for managing BPB in rice plants.

## Acknowledgment

We thank the financial support provided by University of Putra Malaysia UPM. We also thank the Iraqi Ministry of Agriculture represented by the Karbala Agriculture Directorate.

## Data Availability

Data will be available upon request.

